# ORT: A workflow linking genome-scale metabolic models with reactive transport codes

**DOI:** 10.1101/2021.03.02.433463

**Authors:** Rebecca L. Rubinstein, Mikayla A. Borton, Haiyan Zhou, Michael Shaffer, David W. Hoyt, James Stegen, Christopher S. Henry, Kelly C. Wrighton, Roelof Versteeg

## Abstract

**Motivation:** Nutrient and contaminant behavior in the subsurface are governed by multiple coupled hydrobiogeochemical processes which occur across different temporal and spatial scales. Accurate description of macroscopic system behavior requires accounting for the effects of microscopic and especially microbial processes. Microbial processes mediate precipitation and dissolution and change aqueous geochemistry, all of which impacts macroscopic system behavior. As ‘omics data describing microbial processes is increasingly affordable and available, novel methods for using this data quickly and effectively for improved ecosystem models are needed.

**Results:** We propose a workflow (‘Omics to Reactive Transport – ORT) for utilizing metagenomic and environmental data to describe the effect of microbiological processes in macroscopic reactive transport models. This workflow utilizes and couples two open-source software packages: KBase (a software platform for systems biology) and PFLOTRAN (a reactive transport modeling code). We describe the architecture of ORT and demonstrate an implementation using metagenomic and geochemical data from a river system. Our demonstration uses microbiological drivers of nitrification and denitrification to predict nitrogen cycling patterns which agree with those provided with generalized stoichiometries. While our example uses data from a single measurement, our workflow can be applied to spatiotemporal metagenomic datasets to allow for iterative coupling between KBASE and PFLOTRAN.

**Availability and Implementation:** Interactive models available at https://pflotranmodeling.paf.subsurfaceinsights.com/pflotran-simple-model/. Microbiological data available at NCBI via BioProject ID PRJNA576070. ORT Python code available at https://github.com/subsurfaceinsights/ort-kbase-to-pflotran. KBase narrative available at https://narrative.kbase.us/narrative/71260 or static narrative (no login required) at https://kbase.us/n/71260/258

**Supplementary information:** Supplementary data are available online.

## 1 Introduction

The critical zone (CZ) – the area between the top of the forest canopy and the bottom of the groundwater table is essential in sustaining life (Guo and Lin, 2016). Being able to understand and predict critical zone function is essential for both scientific and operational purposes. This understanding and prediction requires the accurate representation of key hydrobiogeochemical ecosystem processes which occur and interact in the critical zone. These ecosystem processes operate at different scales and have different drivers, but at the same time are tightly interconnected. For instance, while hydrological processes control the movement of water at macroscopic scales and are driven by groundwater table gradients, precipitation, and evapotranspiration whereas microbiological processes (Anantharaman *et al*., 2016; Long *et al*., 2016) occur at the microbe scale and are driven by microbial populations, soil properties, aqueous geochemistry, and temperature. However, processes at these two scales influence one another in many ways.

One well-established approach to obtaining an understanding of critical zone behavior is through the use of reactive transport models (RTM) which can simulate coupled chemistry, flow, and transport in hydrobiogeochemical systems. There are a variety of reactive transport codes (see (C. I. Steefel *et al*., 2015) for a review). These models are generally continuum scale models which represent subsurface properties on grids, with grid volumes on the order of cubic meters. As the earth is a porous media with grains and pores, such continuum scale models obviously do not capture pore scale properties and dynamics. One fundamental challenge in numerical modeling is thus how to link and couple processes and properties which happen at different scales (Battiato *et al*., 2011; Chu *et al*., 2012, 2013; Carl I. Steefel *et al*., 2015). Such linking is especially required between macroscopic system behavior and microbial processes which change aqueous geochemistry and mediate precipitation and dissolution.

With continued decrease in ‘omics data analysis costs, one promising approach for this linking is through the incorporation of site-specific microbiological data into RTM to represent microbe-catalyzed biogeochemical more accurately than using generalized stoichiometries. The feasibility of using the results of microbiological data analysis to parameterize RTM has been shown previously. For instance, Scheibe et al. demonstrated the linking of genome scale models with a reactive transport code (in their case, HYDROGEOCHEM) to improve incorporation of microbiological processes on in situ uranium bioremediation (Scheibe *et al*., 2009). Specifically, they used a genome scale model of *Geobacter sulfurreducens* to populate a lookup table spanning reasonable expected ranges for all combinations of three key system parameters. This was then used to predict the effects of varying concentrations of three key growth factors (acetate, Fe(III), and ammonium) on reduction of uranium (VI) at a systems level. More recently, Song et al. developed an enzyme-based approach for simulating microbial reaction kinetics which captured the overall behavior of a consortium rather than rely on individual taxa within the community and coupled it with reactive transport simulations using PFLOTRAN’s Reaction Sandbox (Song and Liu, 2015; Song *et al*., 2017; Hammond *et al*., 2017). This approach is based on a mechanistic understanding of microbial processes and thus can more accurately predict microbial response to perturbations. However, this approach substantial experimental data, such as enzyme concentrations and kinetics data, as well as advanced microbiological knowledge to implement.

These previous efforts have demonstrated the value and feasibility of accurately representing microbial processes in RTM. This in turn opens up the potential to integrate microbiological, geochemical, and physical subsurface properties and processes to predict ecosystem behavior and response (Fig. 1).

**Fig. 1.**
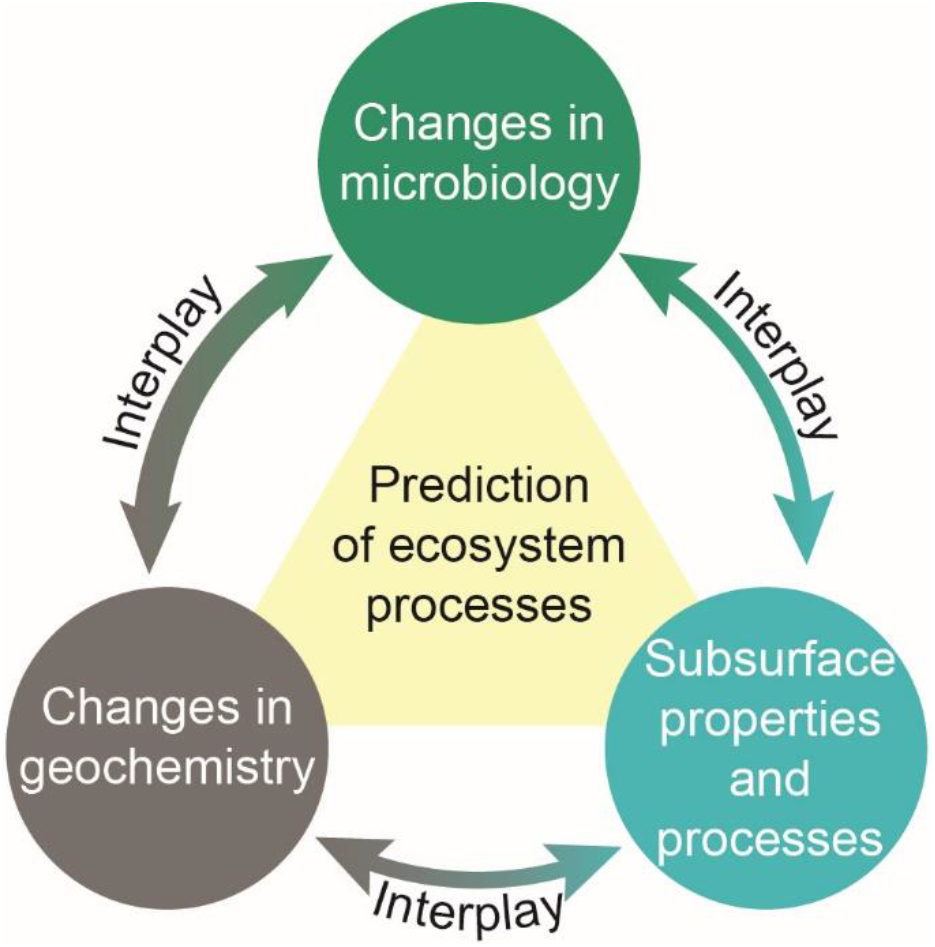
Accurately capturing the interplay between microbiology, geochemistry, and physical subsurface properties and processes is critical to understanding and predicting ecosystem processes.

However, each of the approaches described above requires substantial manual effort to implement for a single site, which makes them challenging to scale. The challenge of scaling these approaches limits the ability to rapidly develop models obtain the associated understanding for many sites. An alternative approach (proposed and demonstrated here) is to an approach which allows for automation.

Specifically, in our workflow we use KBase (a cloud-based software platform for systems biology (Arkin *et al*., 2018)), to automatically generate draft metabolic models from annotated metagenome assembled genomes (MAGs) extracted from environmental samples. These metabolic models can be used (still in KBase) to perform flux balance analysis (FBA) on different media compositions. These media compositions are informed by metabolomics and other site-specific chemistry data. The output of the FBA can be used in reactive transport models (such as the reactive transport model PFLOTRAN (Mills *et al*., 2009; Hammond and Lichtner, 2010; Gardner *et al*., 2015)), which we use in this work. PFLOTRAN is an open source, massively parallel reactive transport code which supports multi-phase (e.g. aqueous, gaseous), multi-component (multiple chemical species), and multi-scale (e.g. pore or macroscale) simulation of contaminant transport in porous media, as well as includes a basic implementation of microbial reactions modeled by Monod kinetics. One major benefit of PFLOTRAN is that users can implement custom reactions or kinetics through the Reaction Sandbox (Hammond, 2017). Our workflow, called ‘Omics to Reactive Transport (ORT) (Fig. 2), thus captures both microbial metabolisms based on environmental samples (using KBase) and macro-scale hydrologic and geochemical processes (using PFLOTRAN). In the remainder of this paper we present the concept and implementation of this workflow.

**Fig. 2.**
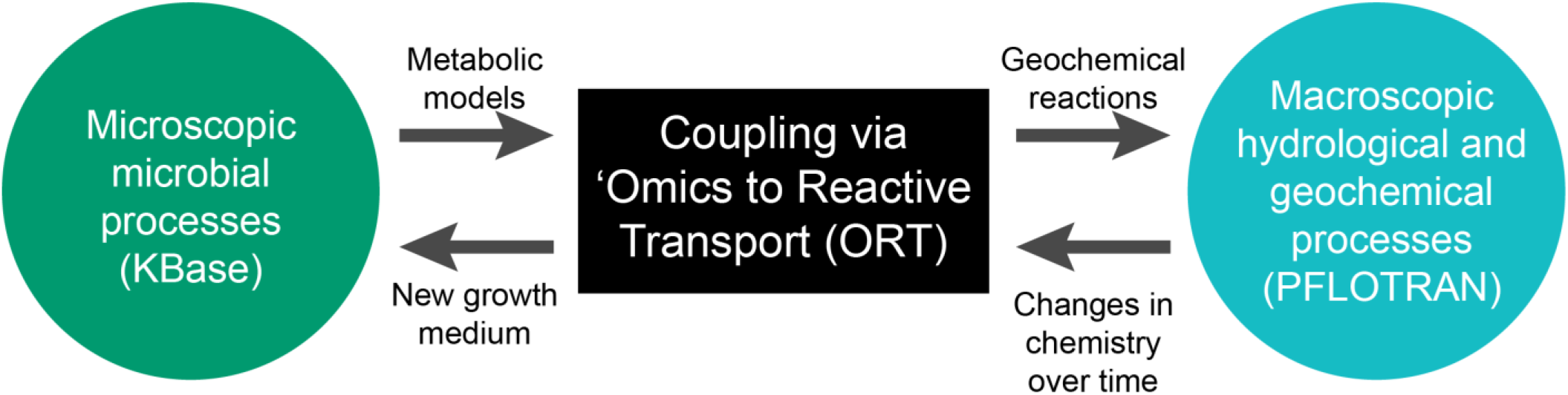
Omics to Reactive Transport (ORT) workflow couples microbe-scale and macroscale processes using the outputs of KBase and PFLOTRAN as inputs for each other.

In addition to the scientific value of this workflow, we want to highlight three operational attributes of interest. First, this workflow can be mostly automated, offering the potential of rapidly generating reactive transport models from microbiological data (detailed in Section 3) with a minimum of manual labor. Second, the resulting models can easily be shared and made accessible to other groups. For instance, we have provided two of the models we generated through a user-friendly web interface which allows end-users to interact with these models. Third, while in this paper we do not include results for this, our workflow lends itself well to an iterative approach. Specifically, it is well known that microbial processes will result in changes in geochemistry, which in turn will influence the microbial processes. In addition to this, macroscopically driven changes in saturation, temperature, and chemistry (e.g., resulting from stage-driven surface water/ground water interaction) will also influence microbial processes. The approach described here can be executed in an iterative manner to capture this two-way coupling between microscopic and macroscopic processes.

## 2 System & Methods

### 2.1 Workflow Concept

The ORT workflow was designed with automation in mind. Specifically, it is designed in a modular manner with a well-defined start and end points and inputs and outputs, with each component being fully automatable (Fig. 3). The inputs to this workflow are annotated genomes, environmental chemistry, and a PFLOTRAN model template which incorporates physical site data. This template (which would be customized to the specific site) would be something like “0D batch reactor” or “2D model of unsaturated soil” (where “nD” indicates the number of spatial dimensions accounted for in the model grid).

**Fig. 3.**
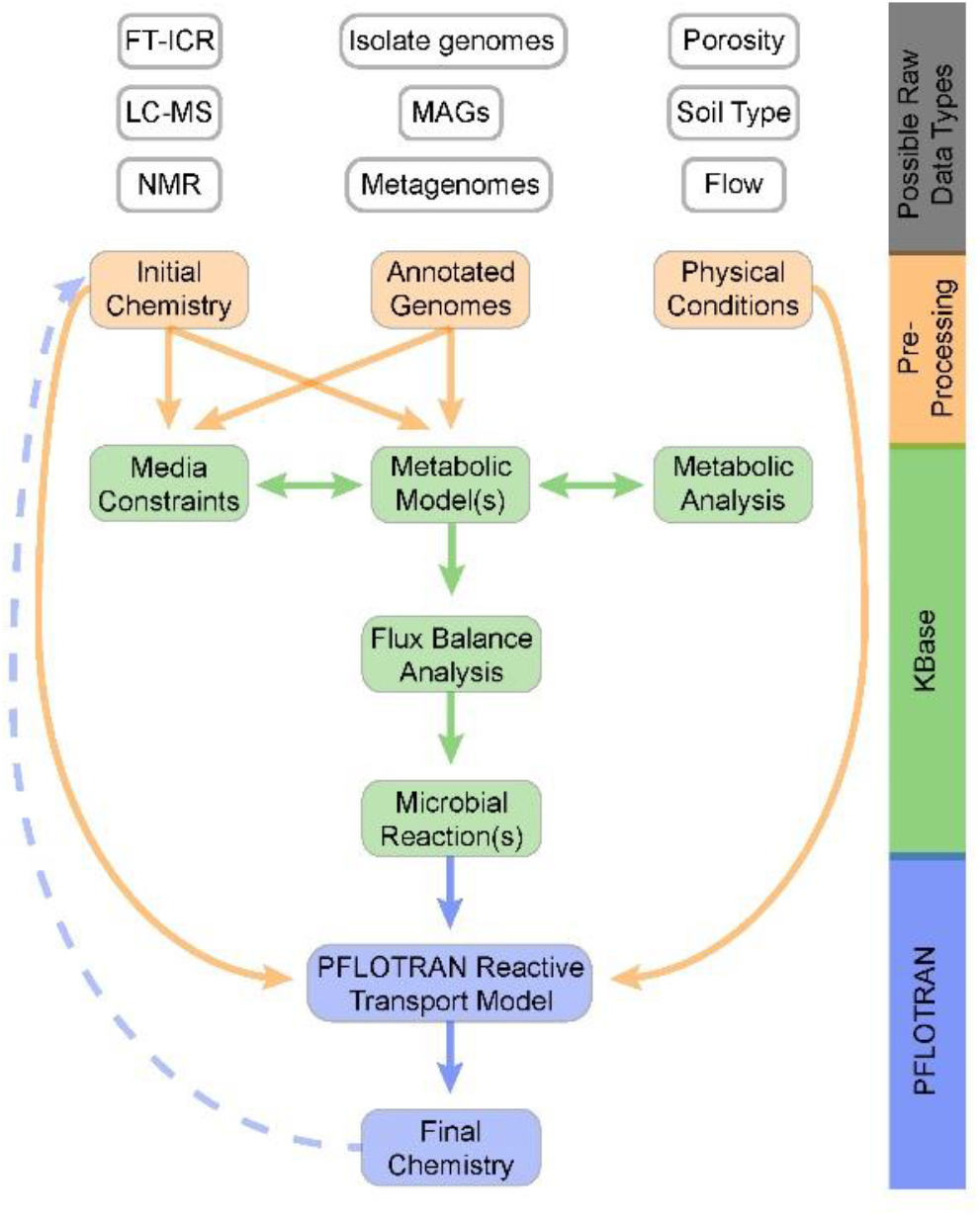
Flowchart of the ORT workflow where orange boxes are workflow inputs based on site characterization which are pre-processed before use, green boxes are metabolic modeling steps carried out in KBase, and blue show the resulting RTM. The horizontally-aligned boxes and arrows in the KBase workflow represent robust curation steps (discussed in Section 4), and the dashed arrow indicates the iteration path wherein the PFLOTRAN-simulated chemistry is used as a new media condition in KBase.

**In** this workflow we import annotated genomes (Shaffer et al., 2020) and site chemistry (e.g., available carbon sources, electron acceptors, and micronutrients based on metabolome and any other chemical analysis at the site, synthesized into a KBase media recipe) into KBase, then use KBase apps to generate the overall reactions. Next, these reactions, as well as the site chemistry, physical site data, and model template are used to build the actual PFLOTRAN model. This model can then be used to simulate macroscopic system behavior.

In the iterative implementation, the PFLOTRAN model simulates changes in physical and chemical conditions in space and time. We can then use the simulated chemistry as a new media composition to be used by the KBase part of the workflow and repeat the process to generate and retrieve new resulting overall reactions and substitute them into the PFLOTRAN input file.

### 2.2 Workflow Implementation

The ORT workflow consists of Python scripts, KBase narratives, and PFLTORAN models. In our implementation of ORT, KBase apps import genomes and chemical data into KBase and use these as inputs for KBase metabolic modeling apps (process described in detail in Sections 2.4 and 2.5). After the completion of the KBase part of the workflow, the KBase API (application programming interface) programmatically exports the KBase-predicted exchange fluxes from KBase. These fluxes are translated by our Python script into an overall reaction string that describes chemical uptake and secretion from each modeled organism, written in PFLOTRAN-compatible naming conventions. The flux values are used as the stoichiometric coefficients for the corresponding chemicals in the overall reaction used in PFLOTRAN, with positive fluxes indicating reactants and negative fluxes indicating products. The summation of exchange fluxes is not a chemical reaction in the traditional sense, but represents the chemical species removed from and added to the system as a result of the microbial metabolism. Thus, this “pseudo-reaction” provides the information needed by PFLOTRAN to simulate the resulting changes in chemical concentrations.

The ORT Python script outputs a \**.txt* file with the reaction strings and yield terms for use in the MICROBIAL_REACTION card in PFLOTRAN as well as a set of \**.dat* files which contain compound names and details which need to be added to the PFLOTRAN geochemical database (formatted for compatibility with the database). This step can either be done programmatically or manually by substituting the content of these text files into a PFLOTRAN model input file (known as an infile). This script bridges the disconnect between KBase and PFLOTRAN illustrated in Fig.2.

### 2.3 Test Case

#### 2.3.1 System Description

To evaluate the performance of our workflow, we used environmental samples from a hyporheic zone in the Columbia River. In these zones, biological nitrogen cycling is known to occur (Triska *et al*., 1993; Zheng *et al*., 2016). Biological nitrification and denitrification is a classic, well-understood, and extensively studied system. We can calculate and compare models which use traditional (textbook) stoichiometries for nitrification and denitrification to the model generated from our workflow. In the remainder of this paper, we refer to these two models as the “literature based model” and “genome derived model”.

Nitrification is traditionally split into two sub-processes, ammonium oxidation (NH_4_ → NO_2_) and nitrite oxidation (NO_2_ → NO_3_), while denitrification is often represented as a complete process (NO_3_ → N_2_), though in reality it is several sequential reactions. Within KBase, we could implement separate models for each step for which genomes are available, but for comparison to the traditional model we used a single model for complete denitrification in this test case. The overall reactions used for the nitrification step were based on experimentally-determined stoichiometries (Liu and Wang, 2012) determined by fitting data collected from bench-scale reactors to traditional half-cell reactions (Rittmann and McCarty, 2012), as given by the following reactions:

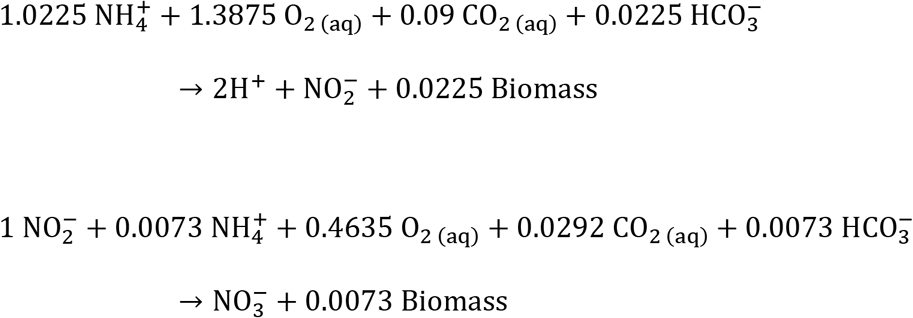

The complete denitrification process stoichiometry was derived from half-cell reactions (Rittmann and McCarty, 2012), scaled to one unit nitrate utilization for comparability with the first two reactions:

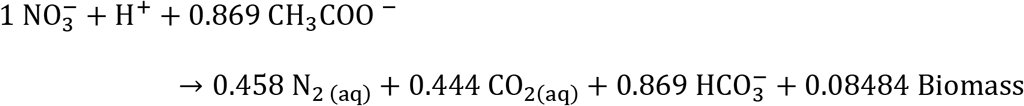

In both cases, the chemical species represented are limited to classical compositions, which in some cases may serve as analogs for a range of compounds. These stoichiometries are not associated with any specific microbes or metabolic pathways, but rather represent the exchange fluxes observed. While this approach is very effective for process design, it does not offer much insight into the microbiology of a system, and may obscure finer-scale dynamics – such as less obvious resources that may become limiting or change how the microbes process available macronutrients, particularly in systems with complex carbon sources.

The rates determined through batch kinetics tests (Liu and Wang, 2012) were used for ammonium oxidation and nitrite oxidation and the denitrification rate was based on rates reported in the literature (Raboni *et al*., 2014). The same rates (shown in Table 1) were used for both the literature-based and genome-based models (described in Section 2.5 - 2.66) in order to directly compare the effects of the different stoichiometries. In future enhancements, we anticipate that reaction rates could be used as tunable parameters to fit these models to system-specific experimental data.

**Table 1.** Baseline reaction rates used in nitrogen-cycling models

| Process | Rate (mol/L·s) |
| --- | --- |
| Ammonium Oxidation | $1.0 \times 10^{-7}$ |
| Nitrite Oxidation | $8.51 \times 10^{-8}$ |
| Nitrate Reduction | $2.34 \times 10^{-8}$ |

#### 2.3.2 Leveraging Existing Multiomics Data

This study made use of multiomics data from previously published work. Sediment was collected and DNA extracted as previously described (Graham et al., 2017). To identify the metabolites available to microorganisms in these river sediments, we performed 1H Nuclear Magnetic Resonance (NMR) spectroscopy on 17 paired sediment pore water samples which also had microbial DA extracted, as described previously (Tfaily *et al*., 2019). Briefly, sediment samples were mixed with water in a 1:1 ratio and then diluted by 10% (vol/vol) with 5 mM 2,2-dimethyl-2-silapentane-5-sulfonate-d6 as an internal standard. The 1D 1H NMR spectra of all samples were processed, assigned, and analyzed using Chenomx NMR Suite 8.3 with quantification based on spectral intensities relative to the internal standard as described. To obtain a representative bulk summary of the metabolite environment in these sediments, the concentration of 31 of the NMR identified metabolites was averaged across the 17 sediment samples, and this data was used as the chemical data input in our ORT workflow (data available in Supplementary Table S1).

Purified genomic DNA was sent to the Joint Genome Institute (JGI, n=33) under JGI/EMSL proposal 1781 and to the Genomics Shared Resource facility at The Ohio State University (OSU, n=10), producing 43 metagenomes from 34 sediment samples with an average sequencing depth of 3.84 (JGI) 25 Gbp (OSU) per sample. JGI and OSU sequencing was performed as previously described in Graham et al (Graham *et al*., 2018) and Borton et al (Borton *et al*., 2018) respectively. Raw reads were processed, assembled, and binned as outlined in previous publications (Shaffer *et al*., 2020) or via the Wrighton Lab GitHub Page (https://github.com/TheWrightonLab). The genomes are available on NCBI via BioProject ID PRJNA576070.

From the sediments, we obtained metagenome assembled genomes (MAGs) from which we selected four genomes that represented key parts of the nitrogen cycle. For each stage of the cycle, the most complete genomes capable of filling those roles were selected. To represent nitrification, we chose the most complete genome representatives of the ammonium oxidizing archaea classified by GTDB-Tk (version 1.3.0, as of 1-21-21) as a member of the family Nitrososphaeraceae within the genus TA-21 (previously within the Phylum Thaumarchaeota) and nitrite oxidizing bacterial member of the Nitrospiraceae for nitrification. Given that the expression and activity of nitrite reductase encoded in Nitrososphaeraceae (previously Thaumarchaeota) is poorly understood at this time (Kuypers et al., 2018), we did not incorporate the production of nitric oxide by Nitrososphaeraceae, and focused only on nitrite outputs from ammonification. To represent denitrification, we selected two Gammaproteobacterial MAGs, both classified within the family Steroidobacteraceae. Note that neither of these genomes encoded a gene to produce N2 gas, but the reaction to convert nitrous oxide to nitrogen gas was added to the metabolic models during gapfilling (see Section 2.5). We selected only four genomes to maintain the simplicity of this proof of concept, but the approach could incorporate as many as are needed to capture system behavior. Each nitrogen-cycling genome was annotated using DRAM (Distilled and Refined Annotation of Metabolism (Shaffer et al., 2020)) with default parameters. The raw annotations containing an inventory of all database annotations for every gene from each input genome are included in the online Supplementary Materials. These genomes and their annotations were uploaded to KBase (Section 2.5) and were the basis for the KBase-derived model (Section 2.6).

### 2.4 Pre-Processing

Prior to executing the workflow, we need to gather and preprocess data and make several decisions such as selecting a model template. In this section, we describe the data preprocessing steps in generic terms, as the same steps will be required for any system. To begin our workflow, user inputs were organized and prepared, which consisted of three broad steps:

1. Qualitative assessment – to balance model complexity and utility, the system definition phase began with a qualitative description of the system in terms of model type (batch, chemostat, continuously stirred tank reactor, etc.), important processes (such as nitrification or sulfur reduction, depending on the system), and parameters of interest (pH, specific chemical species, etc.) that can guide model development. This step includes evaluating if there is any “missing” data, which might render the model inaccurate or impossible, and would need to be estimated in order to produce a viable system (for example, concentrations of biologically necessary compounds that were not measured). These are identified through a combination of subject matter knowledge and comparison with KBase default media recipes. Note that this does not entail delineation of every process and parameter involved in the system, but rather selection of those important to the specific research or application. The goal of this step is to develop a conceptual model of the system of interest, which may be augmented and refined as needed to accommodate new data. Because many of these models will be similar (e.g. 0D batch models), we can build a library of model templates which can be readily reused.
2. Data Gathering - data describing the site may be drawn from a variety of sources, including direct sampling at the site and public resources such as weather stations or national databases. Biological data could come in the form of annotated genomes or metagenomes collected from the site, or genomes for key microbes as determined using 16S rRNA gene data or literature review could be drawn from public databases. Chemical data could include traditional geochemical analysis as well as metabolomics and metaproteomics to provide a more detailed picture of the chemical profile at the site. Physical data could include temperature, soil porosity, or other parameters of that nature that would be included in the PFLOTRAN input file to produce a more site-specific model.
3. Translation to KBase and PFLOTRAN - the data produced by the various analyses above are not necessarily in formats that may be directly imported to KBase and/or PFLOTRAN. Therefore, the final step in this phase was to translate these data to forms that can be used by the relevant tools (KBase or PFLOTRAN). Aside from managing file formats (see the KBase documentation for details), one major consideration was accounting for any unmeasured chemical species identified in the first step of the preparation phase that needed to be added to the KBase media composition to make it biologically viable or usable by the metabolic models generated in KBase. Additions were limited to chemical species or compounds known (or reasonably expected) to be present and were added in sufficient concentration that they would not be growth-limiting. The primary check for the presence assumption was that the experimental data indicated that the microbes used were both present and involved in nitrogen cycling at that site. We did not investigate the assumption that these compounds were non-limiting, as this is outside the scope of this work.

### 2.5 KBase Metabolic Modeling

Once pre-processing was complete, we can start the ORT workflow. Genomes were uploaded to KBase as paired FASTA and GFF3 text files using the “Import GFF3/FASTA file as Genome from Staging Area” app and then annotated with RASTtk using the “Annotate Microbial Genome” app in KBase. Additional custom annotations from DRAM were uploaded as flat text files using the beta version of “Import Annotations from Staging” app. If using DRAM annotations, preprocessing may be carried out using the provided script at https://github.com/subsurfaceinsights/ort-kbase-to-pflotran. Notably, both RASTtk and DRAM are available as apps in KBase, allowing users to functionally annotate genomes without high memory computational resources. However, note that the DRAM app in KBase differs from the version used in this example narrative (Shaffer *et al*., 2020) as the KBase DRAM app annotates using KOfam instead of KEGG genes and does not currently include EC reaction identifiers, so end results may differ from the included narrative. Chemical data was uploaded as flat text files using the “Import Media file (TSV/Excel) from Staging Area”. The use of pre-processed flat text files as inputs to the workflow significantly simplifies the process compared to using raw data, especially for genomes, and these can be generated automatically using scripts such as the one developed for the DRAM outputs. This first step brought all of our data in the KBase workspace in an integrated manner.

After this step, we used all this data as inputs to the “Build Metabolic Model” app, and the generated models were used in conjunction with the media objects as inputs to the “Run Flux Balance Analysis” (FBA) app. Going forward, we refer to this pairing as “growing a model,” meaning we ran the analysis to determine if biomass growth was possible under the given chemical conditions. The output from the FBA app included the reaction and exchange fluxes for each model grown on the corresponding media.

### 2.6 PFLOTRAN Reactive Transport Modeling

We used our workflow to download the FBA exchange flux values using the KBase API and translate them from KBase objects with ModelSEED (Henry *et al*., 2010) compound IDs to flat text files with reaction strings written using PFLOTRAN naming conventions. We then used either the KBase-derived reaction strings and biomass yield values or the literature-based stoichiometries introduced in Section 2.3 to fill in the MICROBIAL_REACTION card in our 0D model template. All parameters except the reactions and yield terms were held the same for both the literature-based model and the genome-derived model.

## 3 Model behavior and General behaviors and trends

Both models exhibited sequential ammonium and nitrite oxidation followed by nitrate reduction, ultimately producing dissolved nitrogen gas (Fig. 4). Despite using the same reaction rates, inhibition constants, and initial nutrient concentrations, the overall progress of the system is noticeably different. The genome-derived model exhausts the available ammonium within 1.5 hours of the simulation start, while the literature based model does not exhaust ammonium until a little more than 3.5 hours into the simulation. Nitrite concentration peaks earlier and at a lower level for the genome-derived model (~18 μM at approximately 1 hr) than the literature based model (~51 μM slightly before 3 hrs). Similarly, nitrate peaks at approximately 4 μM after 1.5 hrs for the genome-derived model but peaks at 40 μM at the 6 hr mark for the literature based model. In the 6 hour period shown in Fig. 4, the genome derived model has exhausted ammonium, nitrite, and nitrate, while the literature based model is still processing nitrite and nitrate. This variance is expected since we are comparing generic reactions (with generic substrate utilization and biomass production reactions) to site-specific reactions based on the most dominant taxa found at our study site.

**Fig. 4.**
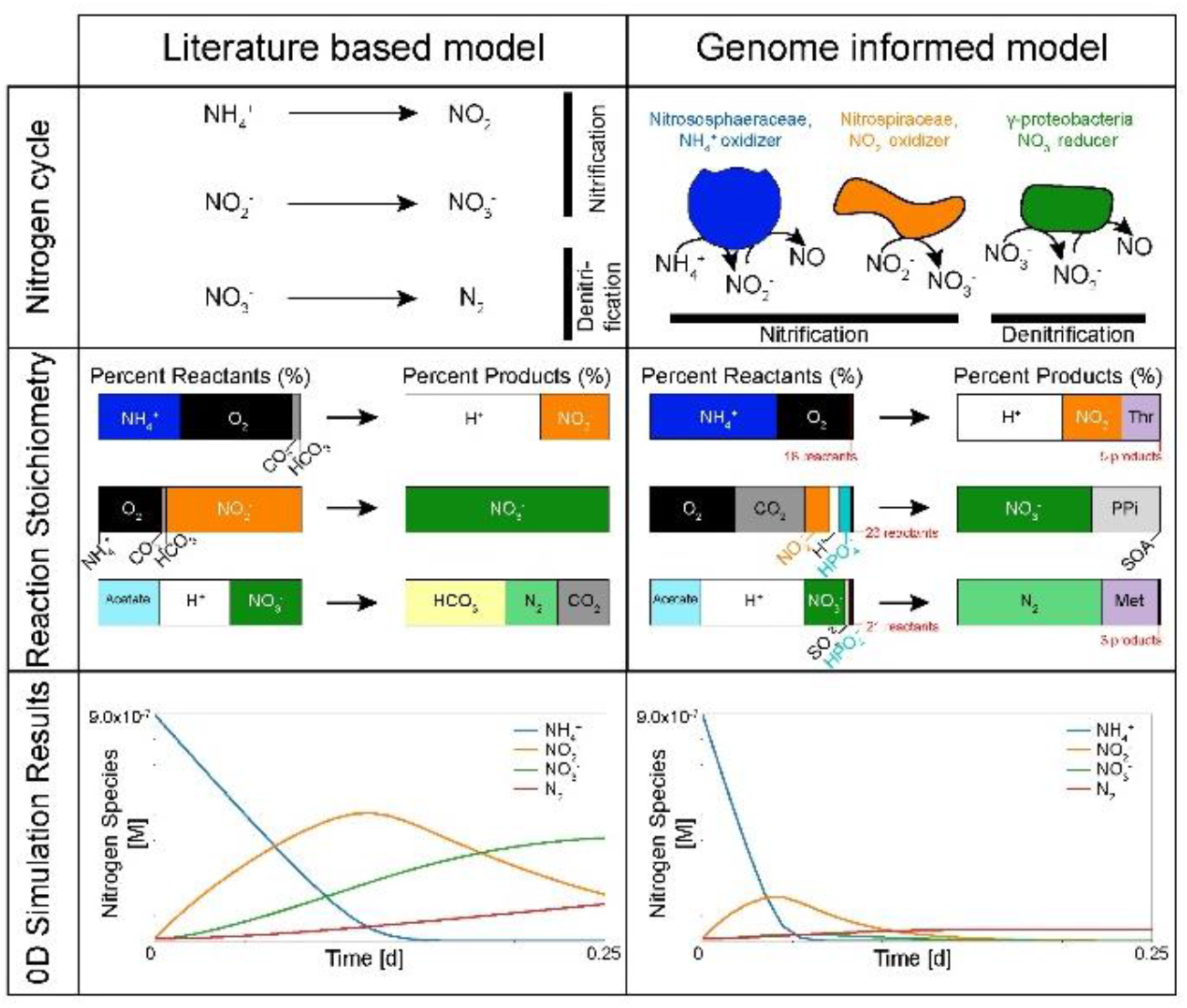
Using our Omics to Reactive Transport (ORT) workflow allows us to not only tailor a model to a specific environmental site and system, but also provides much finer insight into the changes in chemistry driven by microbial processes. The top frame shows the steps captured by the literature-based and genome-informed models respectively. The middle frame shows graphical representations of the two sets of reaction stoichiometries. Abbreviations used in the site-specific model frame are Met for Methionine, Thr for Threonine, and SAO for S-Adenosyl-4-methylthio-2-oxobutanoate, which are compounds predicted by KBase as an output which is not part of standard literature representations. The bottom frame shows the results of using each set of reactions in a 0D PFLOTRAN simulation of nitrogen cycling.

One important difference was that the microbiologically-explicit, genome-based stoichiometry provided much greater detail on the chemistry, particularly with respect to carbon catabolism (Fig. 4 and Fig S1). Specifically, the literature-based models relied entirely on either carbon dioxide (nitrification) or acetate (denitrification), however, because we provided additional carbon compounds detected from our bulk sediment metabolome, the site models used 15 to 23 unique additional carbon sources, such as betaine, leucine, and choline (see Supplementary Table 1). This greater detail allows us to evaluate more precisely the potential chemical drivers or limiters of a system which would be entirely overlooked with traditional representations, which presents the opportunity to probe and improve our conceptual and mechanistic understanding of these systems and individual metabolisms.

Instead of generic bacterial enzymatic reactions, we can determine which site specific bacterial – or archaeal – reactions are drivers in the system. Instead of pre-set stoichiometries, our ‘Omics to Reactive Transport workflow uses chemistry determined based on metabolomics – using these data to describe the initial chemistry rather than generic or simplified chemistry. For example, even with the same rate constants, we can see that the genome-informed model utilizes a higher proportion of ammonium in the first step of nitrification, resulting in more rapid depletion of ammonium in the system and earlier generation of nitrite. As a result, subsequent steps begin earlier, resulting in an overall accelerated process. At the same time, both versions exhibit the expected cycling of ammonium to nitrite to nitrate and finally to nitrogen gas. Since PFLOTRAN relies on user-defined chemistry (as opposed to automatically generating reactions), this allowed us to incorporate more realistic, mechanism-driven reactions.

The genome-based model also allows for greater chemical breadth. The nitrogen cycling reactions are modulated by a wider range of carbon sources. Additionally, the by-products of this carbon and nitrogen metabolism also resulted in more complex chemical outputs in some cases, such as L-Threonine or L-Methionine. These inferred reactions could be further refined by using gene expression data (e.g., metatranscriptomics or metaproteomics data) to calibrate the models (by way of reaction rates, saturation constants, etc.) to a particular set of environmental conditions. Again, this presents an opportunity to test and enhance our understanding of the metabolic processes involved.

Readers can explore and interact with both of these models (without sign-in) through Subsurface Insights’ web-based PFLOTRAN interface at https://pflotranmodeling.paf.subsurfaceinsights.com/pflotran-simple-model/. For the literature-based model, we have made the input concentrations of ammonium, bicarbonate, and acetate accessible to web users using sliders. For the Hanford 300 Area-specific version of the model, we have made accessible the reaction rate for each of the steps modeled. There is no limit to the number of parameters that may be exposed this way, but for the sake of a user-friendly and uncluttered demonstration, we limited our selections to three per model. We selected the parameters we did both because the effects of varying them are significant and to highlight the power and flexibility provided by this approach.

## 4 Discussion

We demonstrated an Omics to Reactive Transport (ORT) workflow for creating site specific reactive transport models that include local chemical and biological content. The ORT workflow was applied to a well-understood system, and the results agree generally with expected behavior in a nitrogen cycling system. We interpret the differences in magnitude and timing to be due to the difference between generic, simplified reactions and metabolism-informed reactions, as KBase-derived stoichiometries made it possible to capture microbial metabolism in much greater detail than conventional approaches allow.

While the model predictions are borne out by comparison to traditional models, we would need extensive new data which currently is not available to comprehensively validate our modeling results. Specifically, we would need high resolution time series data. Such data was not available in this effort, but is a component of ongoing work, and is in general becoming increasingly available as technology improves and cost per sample decreases. Given similar data types, the same workflow could be applied to build and tune a model for other sites.

Much of the future work on this workflow will be focused on enhancing and expanding automation and on making it more robust in several ways. One capability which would be highly beneficial to our workflow is automated metabolic model curation. In our effort, curation was carried out manually using two different approaches: metabolism-based and media-based. The former is labor intensive and requires substantial subject-matter expertise to carry out. The latter is more straightforward and relies on a more general system understanding, but still requires manual iteration to obtain reasonable results. Partially or fully automated model curation will eventually be needed for full automation. This is a topic of active effort by both the KBase core team and other groups, and we will leverage their efforts. Additional work will be in expanding PFLOTRAN models to include processes such as temperature mediated biological processes and material recycling. While these are currently not part of the core PFLOTRAN capabilities, these can be implemented using the PFLOTRAN sandbox.

While previous researchers have demonstrated the feasibility of coupling genome-scale metabolic models with reactive transport simulations, our work is different in some fundamental ways. First, our workflow, lends itself to automation and rapid model generation from ‘omics data. As ‘omics data becomes increasingly affordable, the ability to rapidly translate this data into information on its the implications for macroscopic system behavior will be needed, and our workflow provides a path towards that. Second, our workflow lends itself to easy incorporation of more realistic microbial reaction kinetics (e.g., based on temperature or soil conditions). Third, our workflow lends itself to iteration, which allows us to couple microscopic and macroscopic processes in either direction. Finally, our workflow provides an easy way to couple two powerful and complex software packages which typically are used by scientist in different domains, and allows these scientists a path to generate ‘omics informed reactive transport models.

## Acknowledgements

Tasya Rodzianko, Doug Johnson, and Erek Alper at Subsurface Insights work on the cyberinfrastructure and web interface that was used in this work. Garret Smith, Pengfei Liu, and Lindsey Solden provided additional microbiological expertise and processing. Field sample data was collected by Evan Arntzen, Alex Crump, Brad Fritz, Dave Kennedy, Sarah Fansler, Nate Phillips, Sadie Montgomery, Kyle Parker, and Rob Macklet at Pacific Northwest National Laboratory. Processing of fine sediments was also performed by Ray Clayton and Chris Strickland and cultural support was provided by Doug McFarland and Joy Ferry.

