## Supplemental Material for "ORT: A workflow linking genome-scale metabolic models with reactive transport codes"

**Table S1.** Nutrient concentrations measured in hyporheic zone pore water (site averages)

| Compound | Initial Concentration (μM) |
|---|---|
| Acetate | 131.8 |
| $Ca^{2+}$ | 500 |
| $NH_4^+$ | 0.9 |
| $Mg^{2+}$ | 11 |
| $Fe^{3+}$ | 10.6 |
| $Mn^{2+}$ | 0.9 |
| $HPO_4^{2-}$ | 10 |
| $K^+$ | 3.2 |
| L-Leucine | 0.8 |
| L-Isoleucine | 0.4 |
| Betaine | 0.3 |
| L-Valine | 0.7 |
| Choline | 0.1 |
| Dimethylamine | 0.1 |
| L-Lactate | 2.2 |
| Sucrose | 9.8 |
| Trehalose | 5.7 |
| Melitose | 0.1 |
| 1,2-Propanediol | 0.1 |
| Glycerol | 0.1 |
| D-Glucose | 8.9 |
| D-Fructose | 2.7 |
| $Fe^{2+}$ | 1.6 |
| LGlutamate | 0.1 |
| L-Alanine | 2.1 |

* Note that bicarbonate, carbon dioxide, oxygen, and sulfate are all assumed to be present but were not measured, so these were added to the initial media in low but not limiting concentrations. In addition, the KBase metabolic modeling requires copper, chloride, cobalt, and zinc ions to be present in order for models to run, so these were added in low but not limiting concentrations.

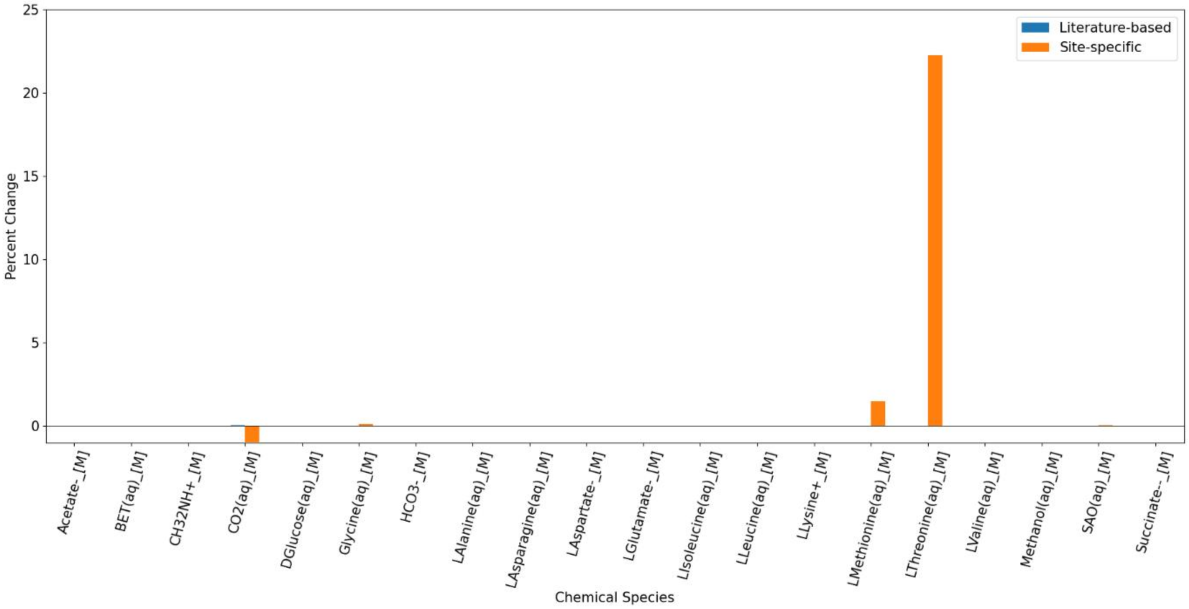


**Figure S1**. Percent change in various carbon sources identified in the bulk soil metabolome. Note that this system was assumed to not be carbon limited, thus the very low percentages. Several of these compounds (Asparagine, Aspartate, Methanol) were produced by one model and consumed by another, and this plot represents only the net change in concentration predicted. The percent change in compounds produced is relative to the baseline initial concentration of $10^{-8}$ mmol used to set up the PFLOTRAN infile.
